# The global β-lactam resistome revealed by comprehensive sequence analysis

**DOI:** 10.1101/2024.03.01.583042

**Authors:** Sevan Gholipour, John Chen, Dongkyu Lee, Nobuhiko Tokuriki

## Abstract

Most antibiotic-resistance genes (ARGs) evolved in environmental microbes long before humanity’s antibiotic breakthrough, and widespread antibiotic use expedited the dissemination of ARGs among clinical pathogens. While widely discussed, the investigation of environmental ARG distributions lacks the scalability and taxonomic information necessary for a comprehensive analysis. Here, we present a global distribution of all five classes of β-lactamases among microbes and environments. We generated a β-lactamase taxonomy-environment map by identifying >113,000 β-lactamases across diverse bacterial phyla and environmental ecosystems. Remarkably abundant, their occurrence is only ∼2.6-fold lower than the essential *recA* gene in various environmental ecosystems, with particularly strong enrichment in wastewater and plant samples. The enrichment in plant samples implies an environment where the arms race of β-lactam producers and resistant bacteria occurred over millions of years. We uncover the origins of clinically relevant β-lactamases (mainly in ɣ-Proteobacteria) and expand beyond the previously suggested wastewater samples in plant, terrestrial, and other aquatic settings.

## Introduction

Long before the discovery of antibiotics by humans, microbes have evolved the capability to withstand antibiotics over the course of their evolutionary history, driven by the arms races between antibiotic producers and those who developed resistance^1^. Unsurprisingly, antibiotic resistance genes (ARGs) have often been observed among microbes living in various environments, including pristine settings unimpacted by human activities^2,3^. The introduction of antibiotics as clinical, veterinary, and agricultural agents generated unprecedented selection pressures on microbes that are related to humans and animals, *e.g.*, pathogenic bacteria^4^. Consequently, many ARGs originally from environmental microbial populations have transferred to pathogens, becoming a major challenge to our healthcare system^5^. Understanding the global distribution of ARGs in various environments and the dynamics of their mobilization from environmental to clinically relevant microbes is critical for devising efficient One Health surveillance systems and strategies to mitigate ARG dissemination^1^. However, the historical research focus on clinically significant ARGs in human-related pathogens has limited our understanding of ARG distribution regarding more diverse microbial species and environments^1^. Factors that remain obscure include the diversity of ARG among microbes, ARG favored taxonomic groups and environments, and the environmental microbes acting as reservoirs for ARG transfer to pathogens. Addressing these questions requires rigorous classification for the sequence profile for each ARG family, followed by a comprehensive study of resistant genes in genomic and metagenomic databases. This approach will reveal the global distribution of ARGs among microbes and diverse environments, allowing the creation of taxonomic-environment maps for ARGs and the identification of reservoirs facilitating ARG transfer to pathogens.

Here, we present a comprehensive, global assessment of the distribution of ARGs by developing tools to overcome previous limitations. We focus on β-lactamases, one of the major ARGs families that significantly contribute to the emergence of multidrug-resistant bacteria, especially gram-negative pathogens^6,7^. These enzymes, with origins dating back 2-3 billion years, have given rise to five distinct β-lactamase classes (A, B1/B2, B3, C, and D) through independent evolutionary events among various microbes^7,8^. To date, over 409 types of β-lactamases have been disseminated among pathogens, all listed in major ARG databases^9,10^.

Numerous prior studies have undertaken major β-lactamase surveys; nevertheless, they mainly focused on highly similar sequences to known β-lactamases with an arbitrary detection threshold^2,11,12^. Consequently, the global distribution of β-lactamases remains hidden. In this study, through comprehensive bioinformatics characterizations with additional experimental validation, we established accurate sequence detection between β-lactamases and closely related homologous non-β-lactamase proteins for each of the five β-lactamase classes. We identified over 113,000 β-lactamase sequences in major public databases (NCBI^13^ and UniProt^14^ and the JGI-IMG metagenomic portals^15^), surpassing the number of sequences in major ARG databases by more than 15-fold. Subsequently, we unveiled the distribution of β-lactamases across various bacterial species and diverse environmental settings, uncovering the origins of clinically relevant β-lactamases that are currently disseminating among pathogenic bacteria through the combination of taxonomic and environmental information.

## Results

### Bioinformatics pipeline for classifying β-lactamase from the databases

A major hurdle in identifying ARGs from sequence data is the lack of understanding of the accurate sequence signature of each ARG family and the sequence borderline between ARGs and closely related non-ARG proteins. To overcome this, we first investigated all β-lactamase families starting from the superfamily-level (**Figure 1a**). Currently, two major β-lactamase families, serine-β-lactamase (SBL) and metallo-β-lactamase (MBL) are known, which are further subclassified into five major classes, class A, B1/B2, B3, C, D^6,7^. Classes A, C, and D belong to SBL and evolved within the penicillin-binding protein-like (PBP-like) superfamily, a protein superfamily that contains enzymes involved in peptidoglycan biosynthesis, such as transpeptidase, DD-carboxypeptidase, and DD-endopeptidase activities^7^. Class B1/B2 and B3 are found in the MBL superfamily, which contains diverse hydrolytic enzymes such as β-lactamases, RNases, phosphonate metabolism, and DNA repair^16^ (**Supplementary Fig. 1a and 2a**).

**Figure 1.**
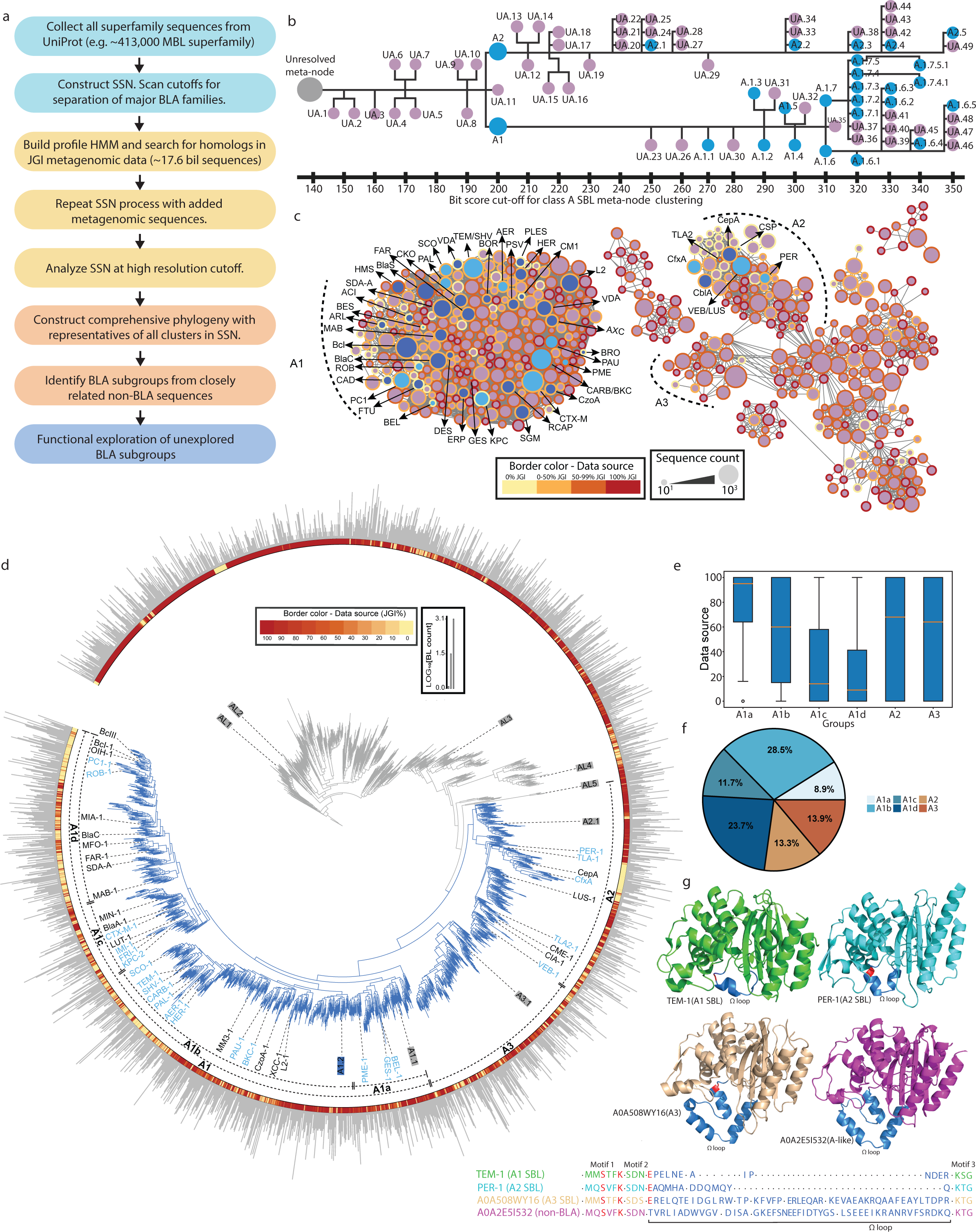
Bioinformatic pipeline for BL identification with class A SBLs as a representative example for the workflow. (a) Workflow to identify BLA subgroups starting from large sequence superfamilies. (b) Example of the bitscore cut-offs at which subgroups of the A family start to emerge as independent clusters. (c) The class A SSN at a bitscore cutoff of 350, the highest resolution cut-off that was deemed necessary according to b. (d) Comprehensive phylogenetic tree of representatives from the A SBLs and neighbouring non-BLA clusters. (e) Box-plot exhibiting the distribution of environemntal sequences in class A clades.(f) Pie-chart exhibiting the size of class A SBL clades. (g) Representative structures and alignment of major class A SBL and non-BLA sequences, hig-hlighting major loop regions.

We first collected all available sequences from the UniProt and NCBI associated with the PBP-like (∼440,000 sequences) and MBL (∼413,000 sequences) superfamilies, and identified sequence subgroups containing known β-lactamase sequences. In general, a protein superfamily with a long evolutionary history forms many separate sequence subgroups corresponding to different families with distinct functionalities^17^. We performed the meta-SSNs (sequence similarity networks) clustering analysis at different “thresholds” (alignment bit scores) to distinguish sequence clusters containing known β-lactamases from other subgroups within each superfamily^18,19^ (**Methods**, **Figure 1b** and **Supplementary Fig. 1c and 2c**). Each β-lactamase class separates out from each superfamily at different clustering thresholds, and ∼10% of the PBP-like superfamily and ∼3.2% of the MBL superfamily are in β-lactamase clusters (**Figure 1b** and **Supplementary Fig. 1b,c and 2b**). Subsequently, for each β-lactamase class (classes A, B1/B2, B3, C, and D), we generated multiple sequence alignments (MSAs) and profile Hidden Markov Models (pHMMs) for the identified sequences, then searched for metagenomic β-lactamase sequences in the Joint Genome Institute (JGI) IMG database^15^ (from 201 projects and 17.6 billion assembled ORFs - **Supplementary File**). All β-lactamase sequences were re-analyzed through SSNs by employing further stringent thresholds for higher resolution of sequences clustering within each class (**Figure 1c and d - Supplementary Fig. 3, 5a, 6a and 7a,c**). We delineated multiple subgroups within each family, which enables us to identify distinct taxonomic or functional β-lactamase subgroups. We also mapped previously characterized β-lactamase sequences, as well as the fraction of sequences each cluster of metagenomic origin (**Figure 1d**). Finally, we cataloged the β-lactamase subgroups from all five classes, establishing the minimum separation threshold of all major subgroups for each class, resulting in 117 B1/B2, 184 B3, 364 A, 78 C, and 211 D subgroups (**Figure 1d - Supplementary Fig. 5a, 6a, and 7a,c**). The unique separations of subgroups within each class highlights the importance of employing different thresholds for each class of β-lactamases. Interestingly, only 124/954 (13%) contain experimentally validated sequences, revealing a substantial reservoir of unexplored β-lactamases. Moreover, we built phylogenetic trees using representative sequences (60% identity for all families) to gain in-depth views of the evolutionary relationship within each class (**Figure 1f, 2 - Supplementary Fig. 5c and 6b**).

**Figure 2.**
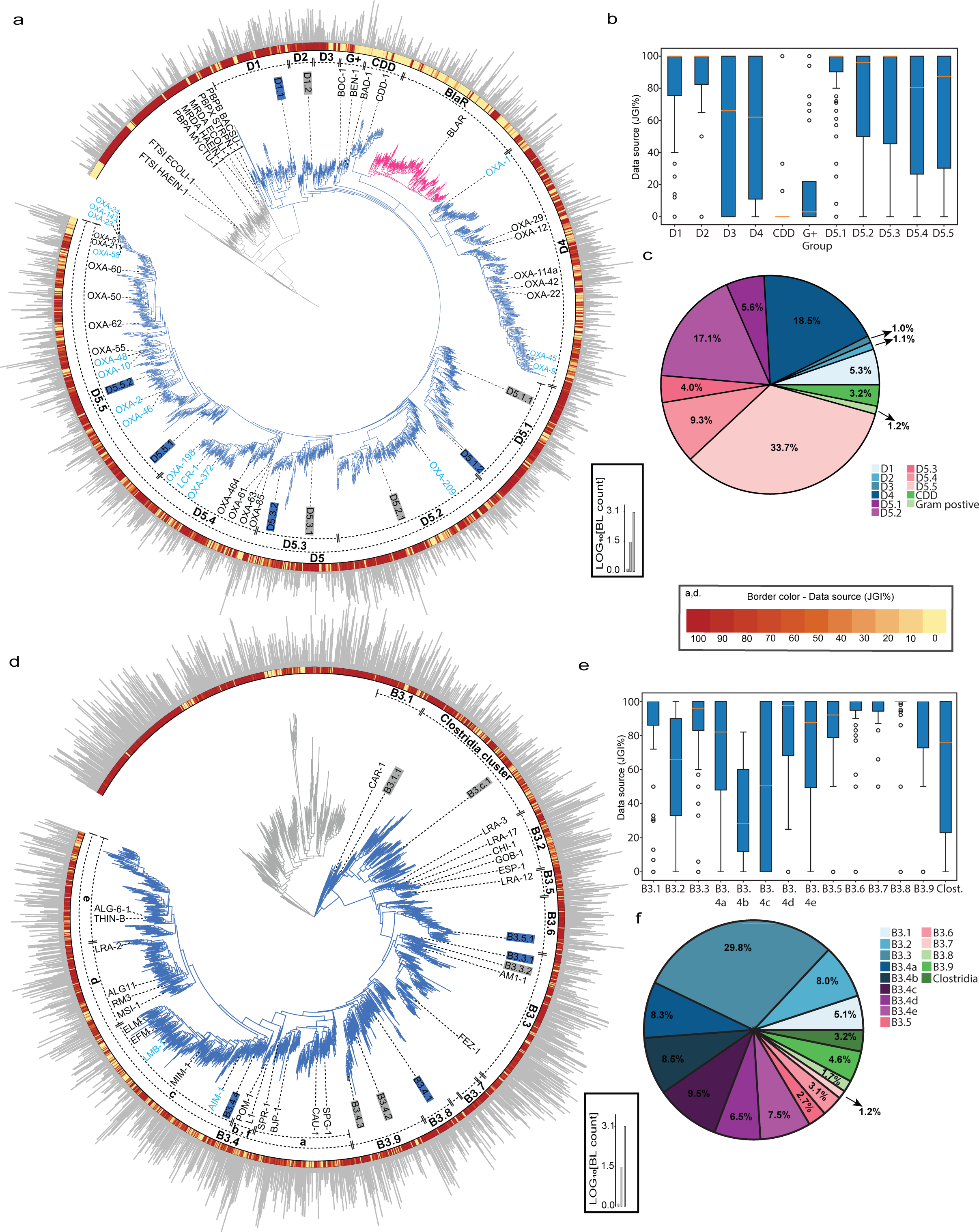
Phylogeny of classes D SBLs and B3 MBLs. Comprehensive phylogenetic tree of representatives from the D (a) and B3 (d) BLAs. In both phylogenies, light blue names represent mobilized sequences, while branch colors indicate the percentage of environmental sequences represented by each phylogeny sequence.The inner dash lines indicate the clade names, while the sequence line color represents the BLA class. The outer bar plot illustrates the number of sequences each node represents. The characterized sequences are visually indicated by the colored box, with active proteins in blue and non-functional sequences in grey. **b** and **e,** Box-plot exhibiting the distribution of environemntal sequences in classes D and B3 clades. **c** and **f,** Pie-chart exhibiting the size of D and B3 clades.

Then, we determined the borderline between β-lactamases and their neighboring non-β-lactamase sequences. Initially, we classified all meta-SSN subgroups in phylogenetic clades containing known β-lactamase sequences as β-lactamase subgroups. Second, we analyzed key sequence and structural motifs of known β-lactamases and compared them to neighboring families (**Supplementary Fig. 3e and 4**). Third, we classified subgroups between known β-lactamase subgroups in phylogenetic clades which displayed the same key motifs as β-lactamase subgroups. Lastly, we performed experimental validation of 40 new genes from unexplored subgroups (**Supplementary Fig. 10e**). To this end, we identified a total of 113,548 β-lactamase sequences (47,332 from NCBI/UniProt, and 66,216 from JGI/IMG) across the five major β-lactamase classes, which is >15-fold the number of sequences currently described in major antibiotic resistance databases^9,10^ (**Supplementary Table 1 and Supplementary Fig. 3**). Moreover, we found a few notable exceptions from conventional β-lactamase clusters. For instance, VarG represents a separate subgroup from B1/B2 classes, and the BlaR transcription regulator subgroup exists within class D. The phylogeny of all five classes are full of unexplored subgroups, highlighting this massive hidden reservoir of β-lactamases.

### Global sequence diversity within each β-lactamase family

In this section, we provide a global view of how each β-lactamase family is distributed. Fully detailed descriptions for each class, including pathogen-related β-lactamases, are in **Supplementary file 1**. Generally, sequences from NCBI/UniProt predominantly showcase β-lactamases from isolated microbes, while metagenomic datasets primarily contain β-lactamases originating from unexplored environmental microbes; there is only ∼3-5% overlap for β-lactamases between these datasets (**Supplementary Table 1**).

In class A SBLs, we classified three primary clades—A1, A2, and A3 (**Figure 1d**). The A1 clade revealed four major subclades (A1a-d), which include major clinically-relevant mobile β-lactamases. While the A1b and A1d are the largest subclades in A1 SBLs with several unknown subgroups, A1a has the highest number of sequences in JGI-IMG environmental samples (**Figure 1e,f**). A2 is the smallest clade, exhibiting a close phylogenetic relationship with the novel A3 subclass, where both clades are major reservoirs of environmental sequences (**Figure 1e**).

For class C SBLs, we identified two major clades, C1 and C2 (**Supplementary Fig. 6b**). C1 is subdivided into C1a-d, with C1b and C1d being most extensive, and C1d being the least studied and containing the majority of environmental C1 sequences (**Supplementary Fig. 6c,d**). C2 encompasses the largest reservoir of class C enzymes, and is divided into C2a-d. C2d represents the largest class C subclade, encompassing all major sequences identified in previous functional metagenomic studies^2^ (**Supplementary Fig. 6d**).

The phylogenetic analysis of class D SBLs revealed the presence of seven significant clades, denoted as D1-5, Gram-positive, and CDD (i.e., *Clostridium difficile* class D) clades^20,21^ (**Figure 2a**). D1-3 are unexplored and prevalent in environmental bacteria, in contrast to the previously established Gram-positive and CDD clades with lower presence in metagenomic samples (**Figure 2b**). D4 and D5 encompass all major clinically significant class D SBLs, constituting the majority of class D sequences (**Figure 2a**). Notably, D5 is prevalent in environmental bacteria and is categorized into five subclades (D5.1 - D5.5), including newly identified subclades such as D5.1 and D5.3 (**Figure 2b and c**). The remaining subclades, D5.2, D5.4, and D5.5, are the primary reservoir of clinically relevant mobile class D carbapenemase (*e.g.*, OXA-10 and OXA-48), with D5.2 and D5.5 being the largest^11^ (**Figure 2c**).

For B1 MBLs, our analysis follows the previous work from Burgland *et al.*, an extensive investigation of B1 MBLs identifying several functional sequences in metagenomic samples^12^. This classification scheme contains five major clades, B1.1–B1.5, with clinically relevant B1 MBLs in B1.1-B1.2 (**Supplementary Fig. 5c**). The B1.3 and B1.4 are notably the largest and remain highly unexplored (**Supplementary Fig. 5e**). In contrast, the B1.5 is the smallest clade, abundant in environmental bacteria (**Supplementary Fig. 5d**), and neighbors two smaller subgroups, B2 and VarG^7,22^. The B2 MBLs are rich in environmental bacteria, whereas VarG is more limited (**Supplementary Fig. 5d,e**).

Lastly, we found 10 major clades within B3 MBLs, the Clostridia and B3.1-9 clades (**Figure 2d**). Notably, the B3.1 and Clostridia clades were previously grouped together, rich in environmental bacteria and featuring only one experimentally validated enzyme (**Figure 2e**). The B3.2 clade is the most extensively investigated, while the B3.3 clade is the second largest, containing major yet unexplored environmental subgroups (**Figure 2e and f**). B3.4 emerges as the largest and most divergent among B3 MBLs, further subdivided into B3.4a-f groups. While B3.4b, c, and f are thoroughly studied subclades with fewer metagenomic sequences, B3.4d-e encompass major unexplored metagenomic subclades (**Figure 2e**). We also uncovered novel B3.5-9 clades widely distributed in environmental bacteria with no experimentally validated sequences.

Altogether, the analyses of all five classes underscores the presence of many underexplored environmental clades. The functional characterization of 40 sequences from these unexplored clades reveals 13 novel functional enzymes across all classes (**Supplementary Fig. 10e**). This includes enzymes with broad spectrum and carbapenemase activity, highlighting the presence of the environmental reservoir of carbapenemase.

### Taxonomic distribution of putative β-lactamase across bacterial phyla

β-lactamase sequences identified in Uniprot/NCBI contains taxonomic information of the host organism, allowing us to investigate the distribution of each β-lactamase class across bacterial domain. Interestingly, each class of β-lactamase has a distinct taxonomic distribution (**Figure 3b**). Overall, MBLs are limited in Gram-positive bacteria compared to SBLs, while both MBLs and SBLs are widely distributed across α-, β-, and γ-Proteobacteria. B1/B2 β-lactamases are predominantly associated with Bacteroidota, with a lower number of sequences in Bacillota and Pseudomonadota. Class B3 MBL sequences are more related to Enterobacterales/Xanthomonadales and Sphingomonadales/Hyphomicrobiales orders in α and ɣ-Proteobacteria, respectively (**Supplementary Fig. 8**). Class A β-lactamases exhibit the most diverse distribution, with significant diversity across Gram-negative and Gram-positive bacteria. Notably, class A is the only group with a diverse reservoir of Actinomycetota-associated β-lactamases, with several members identified in Streptomycetales and Mycobacteriales orders. For class C SBLs, the Pseudomonadota phylum (mainly ɣ-Proteobacteria) harbors the majority of class C SBLs, mainly in Enterobacterales, Pseudomonadales, Hyphomicrobiales, and Burkholderiales orders (**Supplementary Fig. 9**). Class D SBLs share similar taxonomic distribution to class A, except for their low presence in the Actinomycetota phyla^20,21^. Interestingly, the Campylobacterota (previously ɛ-Proteobacteria) phylum is exclusively associated with class D SBLs, harboring several clinical enzymes in the *Campylobacter* spp. (**Supplementary Fig. 9**).

**Figure 3.**
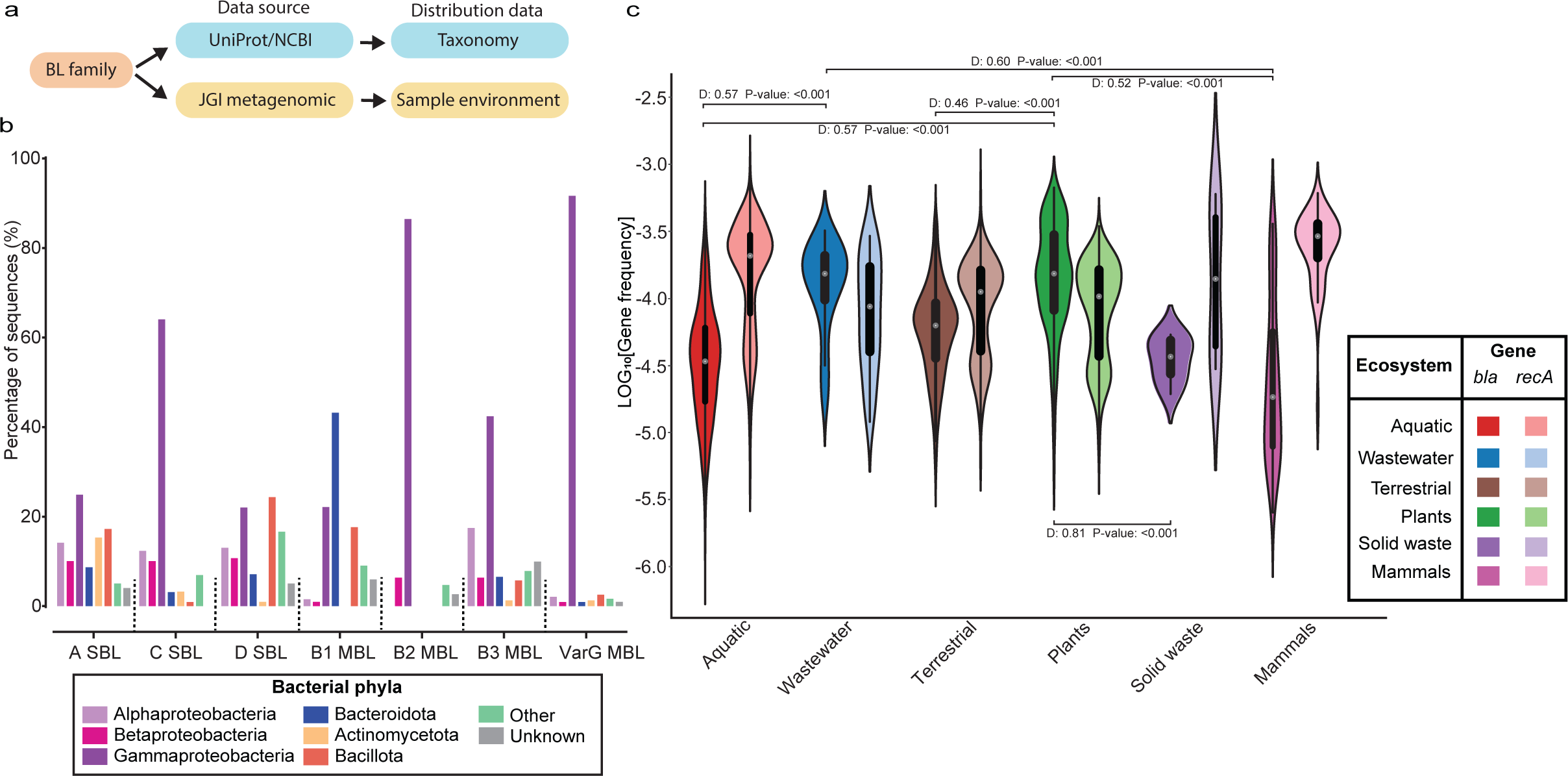
Taxonomic and environmental distribution of β-lactamases. **a**, Schematic of distribution analysis based on data available from the subset of data in each BLA family. **b**, Taxonomic distribution of host bacteria of each BLA family. **c**, Violin plots showing the diversity of β-lactamases and RecA relative frequency in different ecosystems. The summary of the two-sample Kolmogorov-Smirnov test between the distributions are displayed above and below the distributions.

### The distribution of β-lactamases in the environments

Metagenomic data does not often indicate the host organism for an identified gene but contains the sample collection site which identifies the ecosystems where those sequences were isolated. Thus, we analyzed the sample extraction site from the JGI-GOLD database to delineate the environmental distribution of β-lactamases. We calculated the relative frequencies of β-lactamase sequences among all assembled ORFs in each sequencing project, assessing their occurrence in different environments (**Figure 3c**). As a control, we also analyzed the frequency of *recA*, a common single-copy bacterial marker gene that plays a pivotal role in DNA repair and recombination processes^23^. We found unexpectedly high frequencies of β-lactamases across all environmental ecosystems considering that β-lactamases are regarded as non-essential genes. On average, β-lactamases represent ∼4.4 in 10,000 ORFs in the environment samples; only ∼2.6-fold lower than *recA*. This might stem from multiple copies of β-lactamases (*e.g.*, plasmids) in some bacteria, but nevertheless shows that β-lactamases are highly prevalent among environmental microbes. Interestingly, there is significant variation in their relative frequencies depending on the sampled environment. In all ecosystems except for plant-associated and wastewater samples, the relative frequencies of the *recA* marker gene were ∼1.7-to 6-fold higher than those of β-lactamases (**Figure 3c**). On the contrary, in plant-associated and wastewater ecosystems, the relative frequency of β-lactamases exceeded that of *recA* genes by ∼1.6-fold, suggesting a strong enrichment of β-lactamases in these environments^24,25^. The comparison of similar environments in hydrophilic (aquatic *vs*. wastewater) and terrene (terrestrial *vs*. plant) ecosystems further confirmed the enrichment of β-lactamases in wastewater and plant ecosystems (**Supplementary Fig. 10a**) The relative ratios of β-lactamase to *recA* ranged from 1.6 to 3.3 in plant/terrestrial ecosystems and from 1.2 to 6.6 in wastewater/aquatic ecosystems, respectively.

### The taxonomy-environment relationship and origins of clinically relevant β-lactamases

At last, we combined the taxonomy and environmental information to investigate their relationship and the origin of clinically relevant β-lactamases. Our bioinformatics pipeline categorized a total of 954 subgroups spanning all five β-lactamase classes, and among them, 273 subgroups (with a minimum of 25 sequences) included sequences present in both the NCBI/Uniprot and JGI IMG/M databases. Thus, we elucidated the relationship between taxonomy and environmental distribution of β-lactamases by characterizing closely related β-lactamase sequences found among bacterial hosts within the same taxonomic group and/or coexisting in similar environments (**Figure 4**). The majority of subgroups (56%) are exclusive to one phylum. Certain subgroups predominate in one particular ecosystem (*i.e.*, mammals, aquatic environments, plants, or terrestrial habitats), while others are present in multiple settings. Terrestrials and plants were often observed together, while several subgroups were in all three major, aquatic, plants, and terrestrial samples. Considering taxonomy, observing multiple taxonomic classes within a subgroup implies the role of historical horizontal gene transfer (HGT) event(s) among different classes of bacteria that share the same environmental space. This could be detected in Actinomycetota, Bacteroidota, and Proteobacteria phyla. Notably, a significant proportion of HGT takes place predominantly among three major classes of Pseudomonadota, with an occurrence of 63 different class combinations: 15 instances of α-ɣ, 6 of α-β, 24 of β-ɣ, and 18 involving all three. (**Figure 4**).

**Figure 4.**
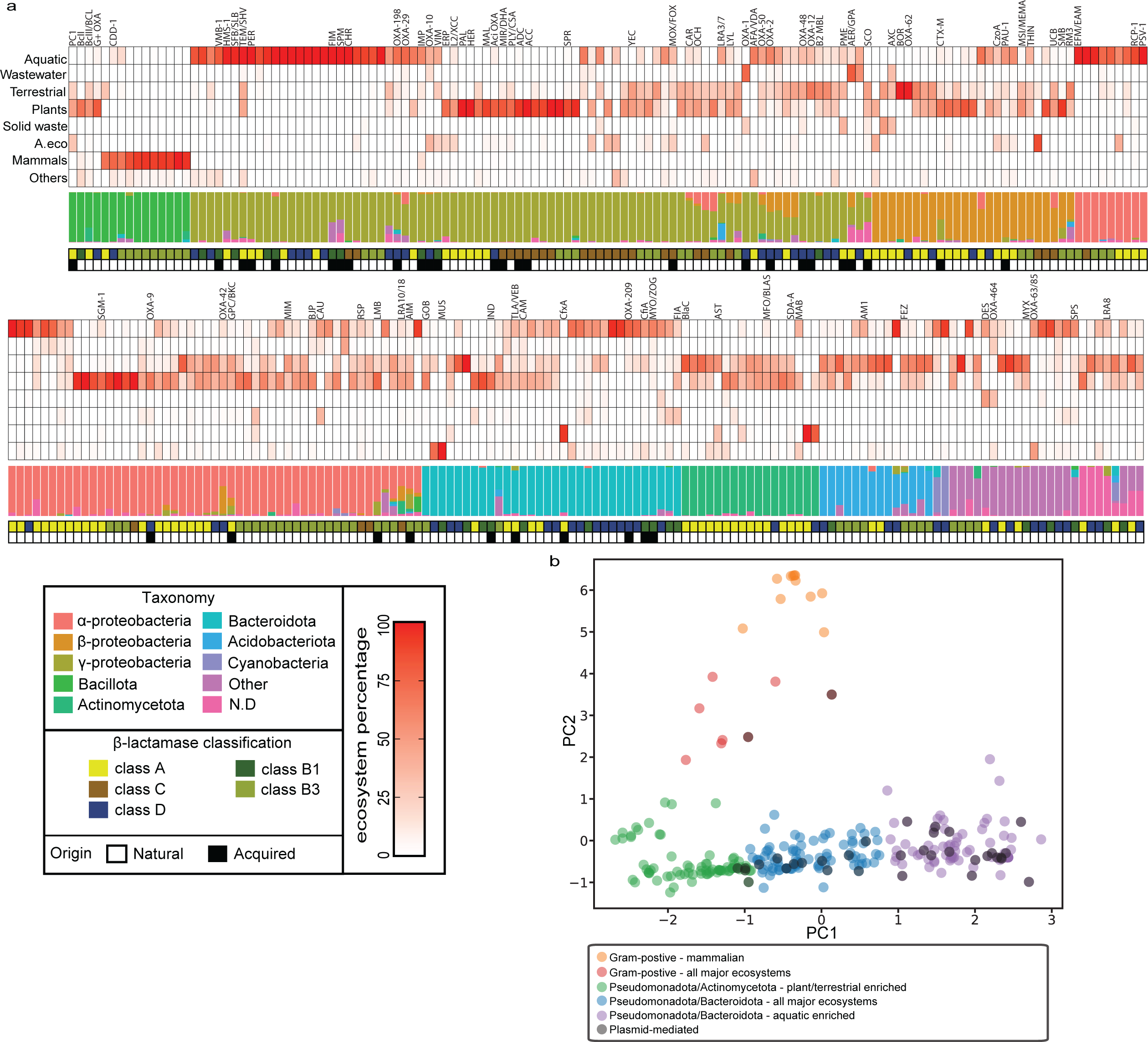
Taxonomic-environmental distribution of β-lactamases subgroups. Heat map illustrates the distribution of environmental sequences in β-lactamase subgroups (more than 25 sequences). Each row in the heat map corresponds to a specific ecosystem, while the columns represent enzyme subgroups. Subgroups containing experimentally validated sequences are labeled above the heat map, while the empty columns indicate unidentified subgroups. The stacked bar plot below indicates the taxonomic composition of the sequence, with their phylum rank corr-esponding to their species IDs. The rows beneath the bar chart specify the class of β-lactamases (above), while the bottom row provides information regarding the origin of the subgroup sequences. **b**, β-lactamase subgroups PCA clustering.The principal component is calculated from the taxonmic/environmental data. The dark subgroups shows subgroups with plasmid-mediated BLA.

Regardless of β-lactamase class, 23 out of 36 clinically mobile β-lactamases (“acquired” in **Figure 4a**) belong to subgroups that are mostly found in ɣ-Proteobacteria, with relatively low distribution in α and β-Proteobacteria and Bacteroidota phyla^11,26^. Some subgroups are found in only aquatic or plant ecosystem, while others are in multiple ecosystems (plant, terrestrial and aquatic) (**Figure 4b)**. The most prominent reservoir for the HGT of β-lactamases into pathogens is ɣ-Proteobacteria found in the aquatic-ecosystem. Additionally, other bacterial phyla and ecosystems host a smaller number of plasmid-mediated subgroups, while those associated with plant ecosystems harbor only a limited number of mobile β-lactamases.

To gain deeper insights into the possible origin of clinically relevant β-lactamases, we further investigated the environmental distribution of sequences closely related to clinical β-lactamases (>90% sequence identities to known β-lactamases). We identified 36 types of clinically significant mobile β-lactamases, including TEM, SHV, CMY, IMP, and OXA-10 types (**Figure 5**). Nevertheless, close homologs of 46 mobile β-lactamases were absent in our environmental dataset, underscoring the need to conduct more extensive metagenomic studies. Interestingly, the preponderance of closely related β-lactamases is discovered within plant-associated samples, with aquatic samples trailing behind. However, β-lactamases closely related to plasmid-borne, mobile β-lactamases showed more diverse patterns. Class C β-lactamases are strongly associated with plant samples, whereas other classes are found in more diverse samples including aquatic and terrestrial samples (Mann-Whitney U test, P-value: 0.0079). Considering the distribution of those closely related β-lactamases may reflect not only the origins of HGT to pathogens but also the results of HGT to plasmids. This suggests the potential for the transfer and dissemination of these enzymes from all major environmental reservoirs. In contrast to plasmid-borne β-lactamases, genomic β-lactamases in pathogenic and environmental bacteria exhibit a predominant presence in samples linked to plants, with limited abundance in aquatic and terrestrial samples.

**Figure 5.**
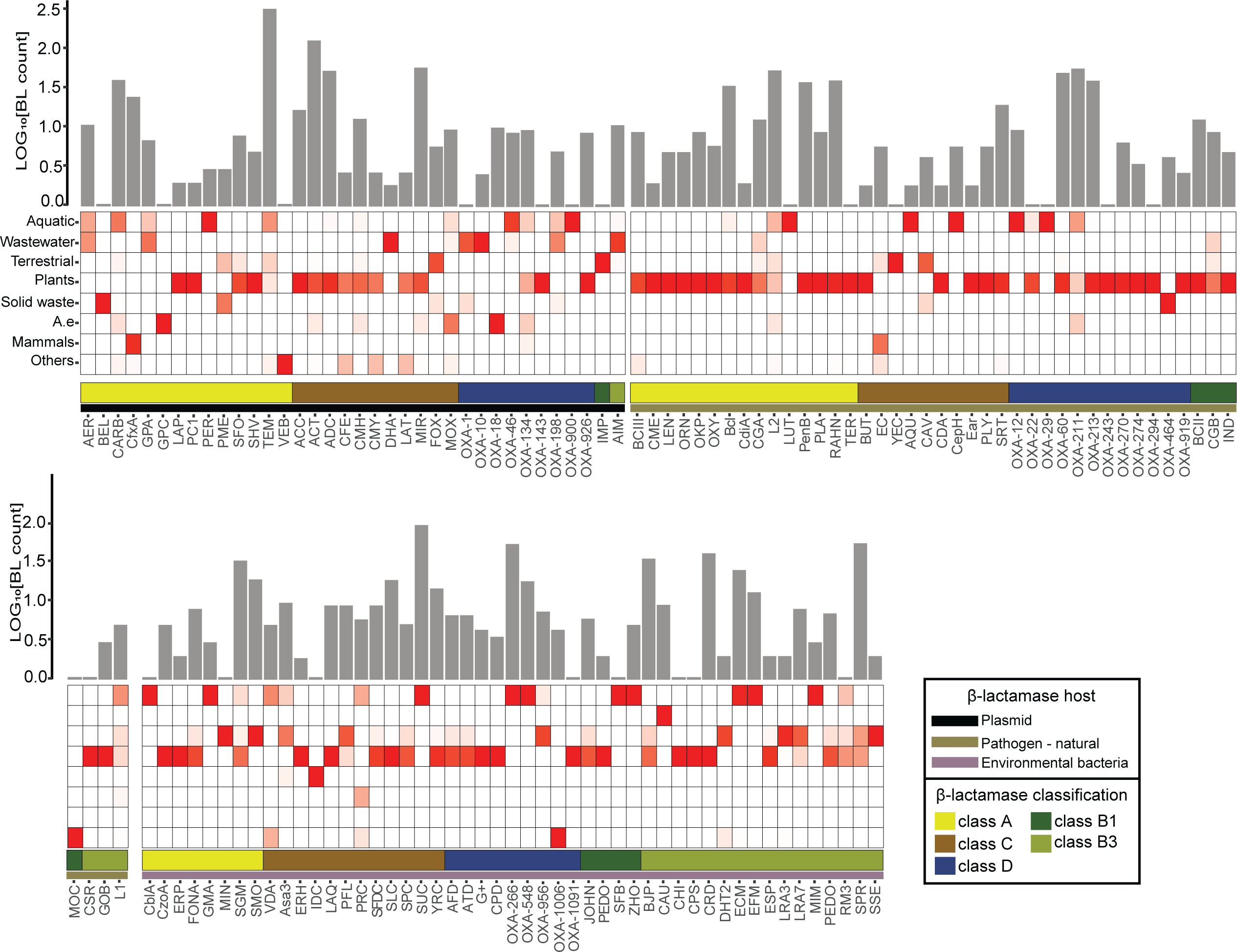
Enviornmental distribution of experimental validated β-lactamases. Heat map illustrates the distribution of experimentally validated β-lactamase variants(sequence identity above 90%). Each row in the heat map corresponds to a specific ecosystem, while the columns represent β-lactamase types. The color strip below the heat map indicates the Ambler classification of β-lactamases. The row below provides information regarding the origin of the subgroup sequences. The bar plot indicates the log scale of the number of each β-lactamase type identified in environmental bacteria.

## Discussion

In this study, we present a global β-lactamase resistome map using our novel comprehensive bioinformatical/experimental workflow. We identify a huge reservoir of β-lactamase sequences across all five classes of β-lactamases, more than 15-fold in sequences listed in the current ARG databases^9,10^. It should be noted that most previous metagenomic studies have utilized arbitrary sequence-search cutoffs for identifying β-lactamases and other ARGs^11,27^ (>75-90% sequences identity to known ARG sequences). We found that such universal and arbitrary cutoffs substantially compromise our ability to identify new ARG sequences. For instance, using sequence identities of >80% or >60% to CARD/BLDB databases only resulted in ∼2% and ∼21% of the metagenomic sequences discovered in this study, respectively (**Supplementary Fig. 10c -d**). This highlights the importance of our approach, which curates β-lactamase sequences from the superfamily level and tests different thresholds to establish rigorous search algorithms. It is also important to note that since each ARG family has substantially different sequence diversity resulting from different evolutionary history, each class of β-lactamase requires a different threshold. Thus, it is essential to perform a comprehensive bioinformatic analysis similar to the methodology employed in this study for all other ARG families to establish rigorous One Health ARG surveillance systems^28^.

We acknowledge the importance of functional characterizations of these newly identified β-lactamase sequences in the future. Each β-lactamase can exhibit substantially different efficiency and specificity toward different classes of β-lactams. Indeed, experimental characterizations of 40 newly identified sequences revealed enzymes capable of degrading carbapenems, which are last-resort antibiotics^29^. Moreover, the lack of large-scale functional information hindered our ability to annotate novel and unexplored β-lactamase subgroups. It is critical to determine whether the β-lactamases transferred to clinically significant pathogens exhibit high efficiency and/or broad specificity against new generations of chemically synthesized β-lactams. Recent developments of multiplex gene synthesis and phenotyping platforms enable the characterization of thousands of proteins, which can unveil the sequence-function maps of many, if not most, environmental β-lactamases^30–32^.

We find that β-lactamases are surprisingly abundant, and widespread in all major bacterial phyla and almost all ecosystems, suggesting their long evolutionary history and the importance of β-lactamases for many bacterial hosts in diverse ecosystems^8^. Generally, only a handful of bacterial species (*e.g.*, *Streptomyces* spp.) and fungi (*e.g.*, *Penicillium* spp. and *Acremonium* spp.) have the capacity to synthesize β-lactam antibiotics^33^. These species are indeed prevalent within various ecosystems, including both aquatic and terrestrial environments, which likely resulted in selective pressure promoting acquisition and retention of β-lactamase genes by numerous microbes within the environment. Further, our study represents the first comprehensive documentation of the potential enrichment of β-lactamases in plant-associated environments. It is known that many *Actinobacteria* spp. have symbiotic relationships with plants enriched in the endophytic compartment in plant rhizosphere^34,35^, with potential to generate β-lactam compounds. This may have created the long arms race between β-lactam producing and resistant microbes, particularly within and around plants over millions of years. It is also plausible that the introduction of fertilizers to soils has an impact on the antibiotic resistance gene (ARG) content, as indicated by previous studies^26^. Moreover, we further confirmed the “human impact” on β-lactamase enrichment, *i.e*., higher frequencies in wastewater, likely linked to the massive production and usage of β-lactams and other chemicals in the last century^24^. Nonetheless, further studies are required to understand the geographical and regional dynamics of β-lactamases dissemination in these environments. The massive reservoir of environmental β-lactamases has enabled numerous β-lactamases to transfer to human pathogens since the introduction of β-lactam antibiotics in clinics. We found that the majority of clinically relevant, plasmid-borne β-lactamases are related to ɣ-proteobacteria. Indeed, this is a major mobilization route of environmental ARGs to clinical pathogens, as most Gram-negative acquired β-lactamases are in ɣ-proteobacteria pathogens^11,26^. While less frequent, the taxonomic distribution of the origin of clinical β-lactamases extends beyond this phylum, suggesting extensive HGT in environmental bacteria. Similarly, the environmental origins of β-lactamases identified within the genomes of pathogenic bacteria display a considerable degree of diversity, while mostly enriched in plant-associated ecosystems. This suggests that plants may serve as a conduit for the movement of ARGs between soil and natural clinical pathogens^26,36^. Interestingly, the taxonomy-environment analysis suggested that the majority of mobile β-lactamases are originated from diverse environments with various levels of aquatic ecosystems presence. Furthermore, aquatic environments allow for more dynamic interactions among microbes, creating opportunities for HGT^37^. In contrast, β-lactamases associated with plant symbionts are confined and remain isolated from other microbes, especially human pathogens. Further studies are required to unveil the detailed HGT dynamics of β-lactamases and other ARGs, and our study offers a substantial foundation to explore and monitor β-lactamases in the environment and clinics.

## Acknowledgments

We thank researchers of the JGI-metagenomic community who permitted to use their data for our work (**Supplementary File**). We also thank members of the Tokuriki lab for the discussions and comments.

## Methods

### Curating sequence dataset

The first step of curating a sequence dataset is to collect data of interest from sequence databases such as Pfam, Interpro, and NCBI. The Pfam and Interpro databases are a collection of protein families defined by structural regions or domains. For SBLs and MBLs, protein sequences of the PBP-like and MBL superfamilies were retrieved from the Pfam and Interpro databases in December 2019 and April 2020 respectively (Pfam-ID: CL0013; Interpro ID: IPR001279). For the NCBI database, β-lactamase sequences from all five classes were extracted using the NCBI BLASTp search, where 10 experimentally validated β-lactamase sequences (BLDB database) from each class were used as queries, and the maximum number of non-redundant hits (20,000 hit with bit scores above 50) were collected. Subsequently, the Pfam and Interpro sequences were combined with the corresponding NCBI search sequences of the corresponding superfamily. The sequences less than 100 amino acids long were removed, and then redundant sequences were reduced by clustering using CD-Hit with a sequence identity threshold of 100%. Next, we extracted a representative set from the combined set of sequences by applying a sequence identity threshold of 60% using CD-Hit^38^. In the last step, we manually added the amino acid sequences of experimentally characterized enzymes from Uniport and BLDB/CARD databases.

Next, 5 HMMs were constructed from the curated datasets from the NCBI and Pfam (**Supplementary Table 1**), representing each major β-lactamase class: i) B1-B2, ii) B3 MBLs iii) A SBLs, iv) C SBLs, and v) D SBLs. The curated sequences were aligned with MUSCLE using the default settings, and HMMs were constructed using the ‘hmm_build’ program of HMMER3^39,40^.

To expand the search to metagenomic data, assembled protein sequences from metagenomic data was extracted from the JGI database in June of 2020. Over 17.1 billion sequences were collected from 201 JGI proposals, covering ∼86% of all assembled genes in the JGI database. We searched the JGI data using the HMM from each of the 5 β-lactamase classes, using the ‘hmm_search’ program, retaining all hits with an inclusive threshold of E-value < 1. The sequence hits with less than 200 amino acids long were removed, a minimum size typical for active β-lactamase with a single domain. Sequences with ambiguous characters (*e.g.*, X, Z) or internal stop codons were removed. To find more closely related homologs, a BLAST search was performed using the genomic β-lactamase sequences from the NCBI and Pfam as queries against the JGI HMM search results, retaining all hits with bit scores above 50. In last step, the non-redundant hits were combined with their corresponding identified sequences in public (NCBI, Interpro, Pfam) databases.

### Calculation and visualization of sequence similarity networks

The PBP-like and MBL representative sequences at 60% identity and all five comprehensive β-lactamase datasets were used as input for the SSN. The all-versus-all BLAST was performed for all sequence datasets^41^. SSNs are generated by clustering sequence hits above a bit score threshold into “meta-nodes”. Each meta-node in a given SSN represents a collection of sequences that have a BLAST bit score above the clustering threshold to at least one other sequence in the same meta-node (https://github.com/johnchen93/MetaSSN) (**Figure 2.1**). In order to search for comprehensive β-lactamase families within the superfamilies, the PBP-like and MBL superfamilies SSNs are clustered at every 10-bit score with a minimum connecting bit score of 10 less than the clustering thresholds, from bit scores of 100 to 300 (e.g., connecting bit score 120 - clustering bit score 130). To perform comprehensive analysis of the β-lactamase SSNs, meta-nodes are constructed at every 10-bit score, starting from of 100 to 350 (100 to 420 for class C). Additionally, experimentally validated MBL (504 B1-B2 and 169 B3) and SBL (160 class A, 61 class C, and 438 class D) sequences from BLDB and CARD databases were used to highlight meta-nodes with characterized β-lactamases. Finally, all networks are visualized in Cytoscape using the organic layout^42^.

### Phylogenetic analysis

All meta-nodes with more than five sequences were used as representatives of each β-lactamase group for phylogenetic analysis. The sequences were retrieved from SSNs at clustering bit scores of 280 (154 B1-B2), 350 (671 class A and 287 B3 meta-nodes), 500 (291 class C meta-nodes), and 300 (233 class D meta-nodes), with the appropriate thresholds for separation being determined from Meta-SSN analysis. Sequences for each meta-node were clustered using CD-hit program at 60% sequence identity threshold. Sequences from each group were aligned using MUSCLE with default settings, and the alignment was curated in Jalview, where the poorly aligned regions and the internal gaps were trimmed according to the structural information^43^. The resulting multiple sequence alignment (MSA) for each class was used for inferring maximum likelihood phylogenies in IQ-TREE 2^44^. Branch supports were assessed through ultrafast bootstrap approximation 2.0 and the approximate likelihood ratio test, both conducted with 5,000 replicates^45^. The sequence evolution model was determined through maximum likelihood using ModelFinder^46^, as implemented in IQ-TREE 2.0 on the representative sequence alignment. All visualizations and analyses of phylogenies were conducted using iTOL^47^.

### Taxonomic and environmental analysis

To obtain taxonomic information from sequences in the NCBI and UniProt databases, the species names were extracted from FASTA sequence IDs. Next, the ete3 toolkit was utilized to convert the species names to NCBI taxids. The obtained taxids were then used to retrieve the full taxonomic ranks for all NCBI and UniProt sequences. For the analysis of environmental distribution, the metadata for all JGI Public Studies/Biosamples was downloaded from the JGI GOLD (Genomes OnLine Database). This information was used to determine the ecological locations of β-lactamase and RecA sequences identified in the JGI database. In order to compare the abundance of these genes across various ecosystems, the relative frequency of *bla* and *recA* in each sample was calculated using Equation (1),

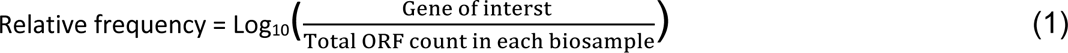

where the putative β-lactamase count is the redundant putative *bla* and *recA* sequences with a length higher than 200 amino acids in each sample.

In the taxonomy-environment analysis, β-lactamase sequences with taxonomic information and the redundant environmental data were gathered from all 273 Meta-node subgroups. These are subgroups with known β-lactamases or a minimum of 25 sequences, selected from the final Meta-SSN clustering thresholds for each class (280 B1, 350 B3 and A, 500 C, and 300 D). These sequences were then compiled for the taxonomic-environmental map.

### Cloning and expression of β-lactamases

To assess the β-lactamase activity of the predicted proteins, 40 proteins were codon optimized for *E. coli* and synthesized using Twist Bioscience gene synthesis service. The synthesized genes were subcloned in a broad-host-range vector, pBTBX-3, along with a modified P5.1 TEM-1 promoter and a pWH1266 origin of replication for *A. baumannii* replication. The plasmids were transformed into *E. coli* strain E. cloni 10G using the chemical transformation protocol.

### Determination of IC50 values

To determine the β-lactamase activity of synthesized genes, single colonies of E. cloni 10G that were transformed with the β-lactamase plasmids were picked and grown overnight at 30 °C in 2.5 mL of LB media with 20 μg/mL of gentamicin for selective resistance. The overnight culture was used at a 1:100 dilution to start a new 5 mL culture of LB media supplemented with 20 μg/mL of gentamicin. The cells were transferred into a 96 well-plate with two-fold increases in the concentration of antibiotic and incubated for 6 h at 37°C. (the range of concentrations screened for each antibiotic was as follows: ceftazidime, 0.25 to 64 μg/mL; meropenem, 0.016 to 1 μg/mL; benzylpenicillin and ampicillin, 2 to 512 μg/mL). The IC50 values were determined at the concentration of antibiotics by which no growth was observed for two replicates.

---

**Supplementary Table 1.**
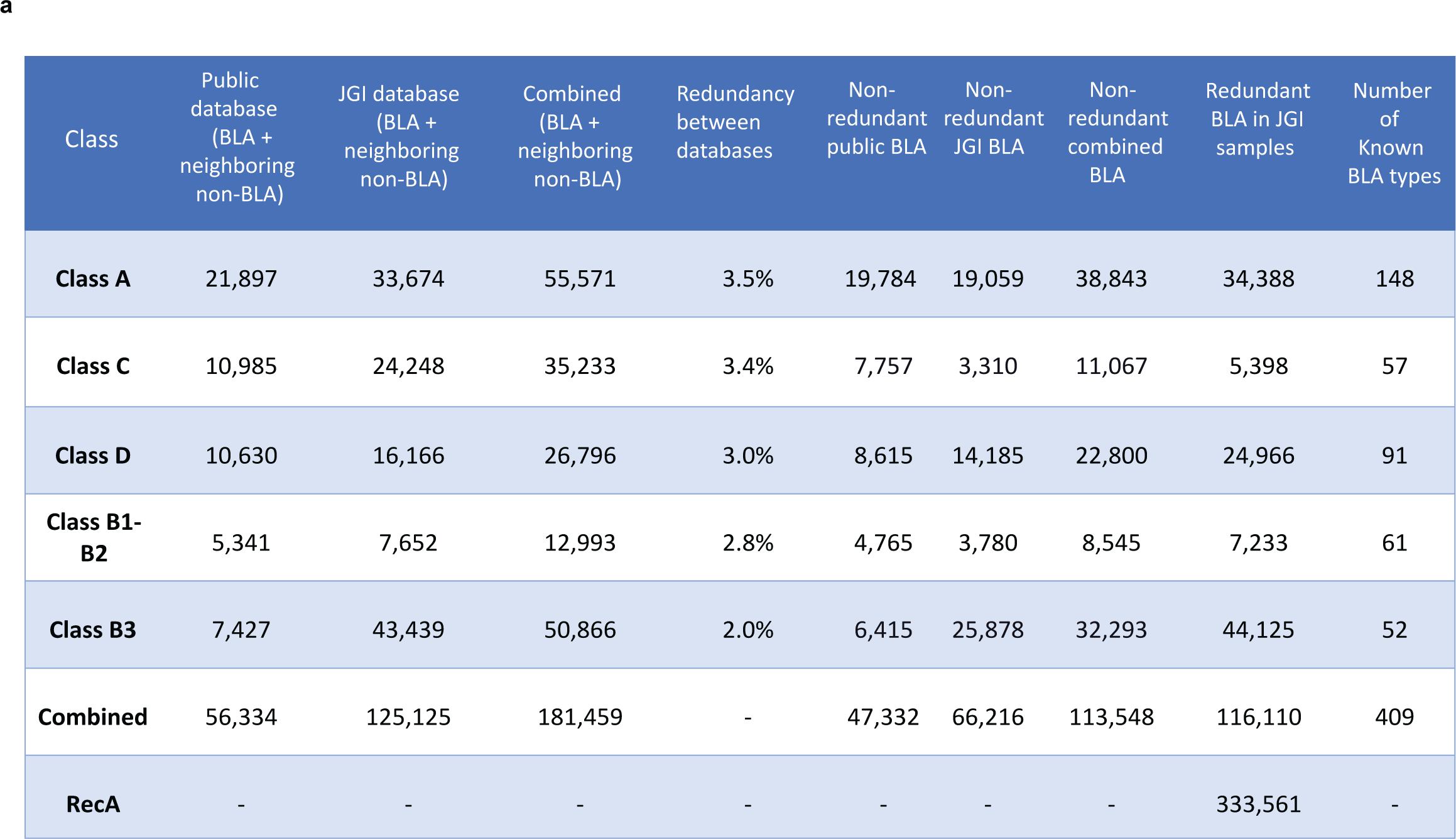
Summary of for putative BLA and RecA sequences in all major databases.

**Supplementary Fig. 1.**
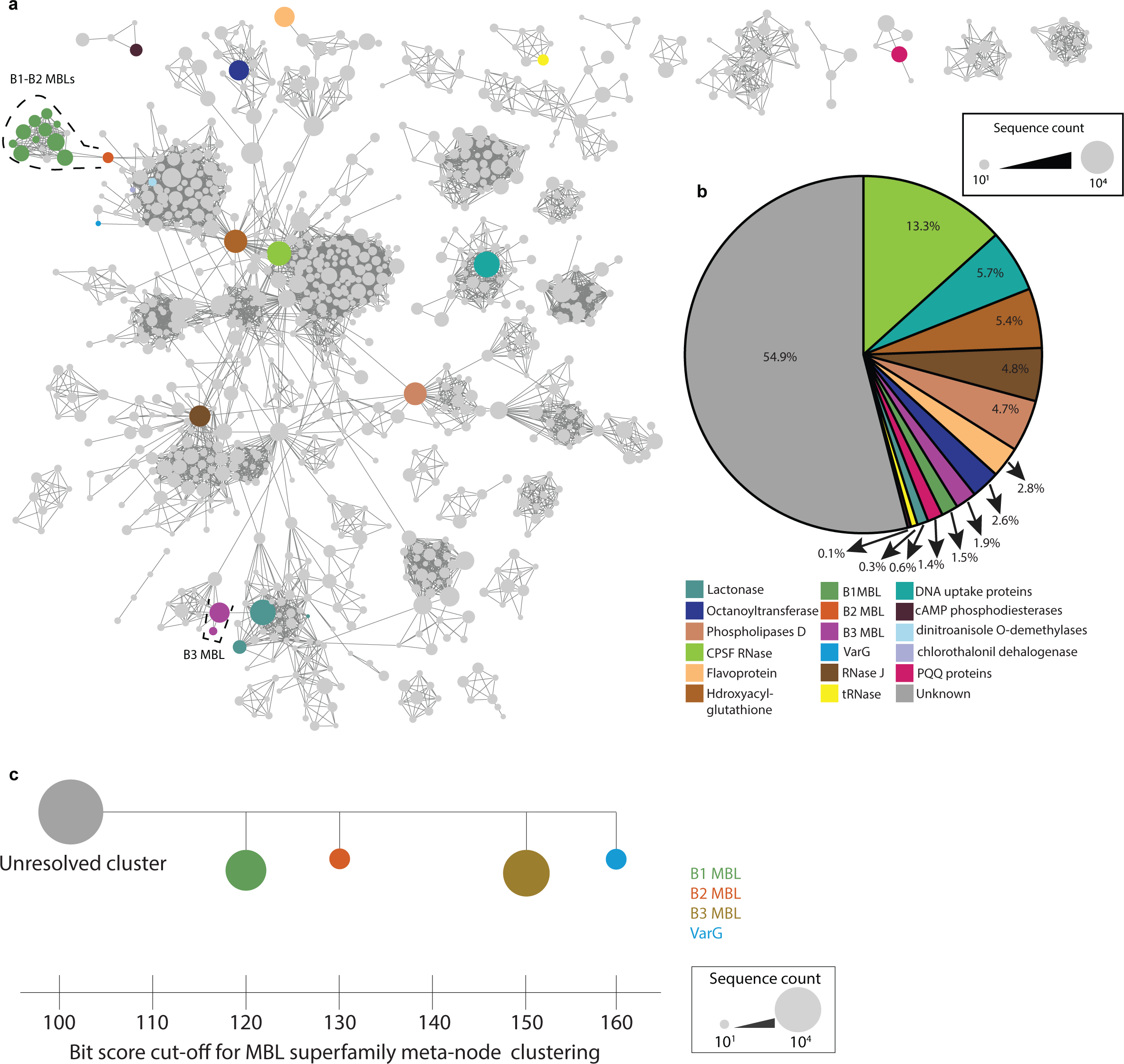
Summary of Metallo-β-lactamases superfamily. Summary of the sequence space for Metallo-β-lactamases (MBL) superfamily. **a**, The sequence relationships among MBL superfamily. Sequence similarity network of MBL superfamily sequences. Nodes indicate groups of β-lactamase sequences sharing BLAST bit scores > 230. Nodes are linked by edges if β-lactamase in two separate nodes share BLAST bit scores > 110. Colored nodes contain at least one sequence found in SwissProt, CARD, and BLDB, while the grey nodes do not have any known or characterized sequences. The size of the nodes indicates the number of sequences for each node in log10 scale. **b**, Distribution of protein families within the MBL superfamily. Categorization of various protein families within the MBL superfamily. The total number of sequences for each subfamily is determined by retrieving sequences from meta-nodes containing experimentally validated sequences and neighbouring connected meta-nodes.**c**, separation of MBL classes in MBL superfamily.Summary of the MBL classes separation bit score cutoff within the MBL superfamily. The bottom axis indicates the SSN clustering threshold defining each class of MBL. The SSN nodes are shown at every 10 bit score, and the exact position of each node does not cover the clustering thresholds in-between bit score intervals.

**Supplementary Fig. 2.**
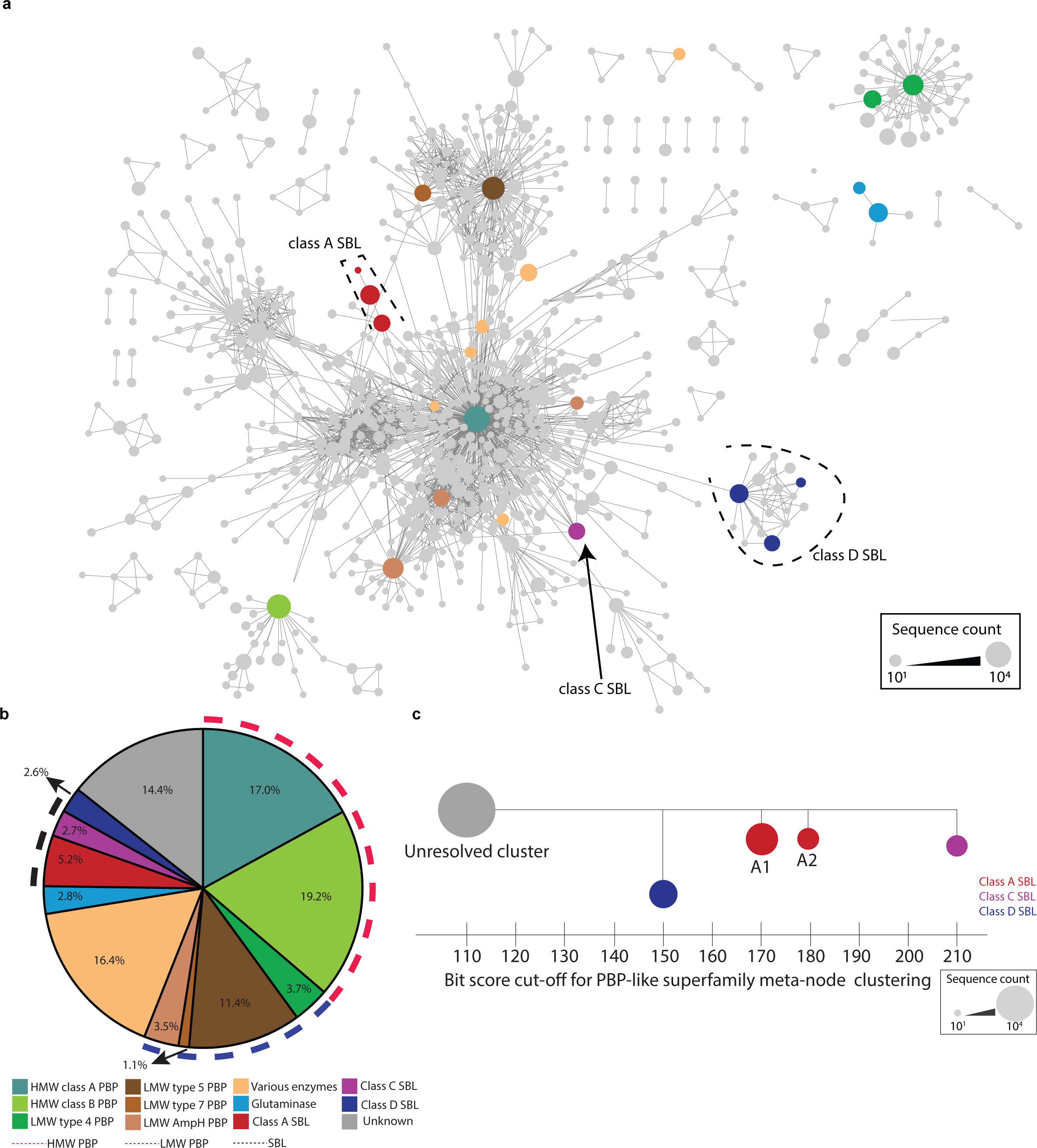
Summary of penicillin binding protein like superfamily. Summary of the sequence space for penicillin binding protein like (PBP-like) superfamily. **a**, The sequence relationships among PBP-like superfamily. Sequence similarity network of PBP-like superfamily sequences. Nodes indicate groups of β-lactamase sequences sharing BLAST bit scores > 210. Nodes are linked by edges if β-lactamase in two separate nodes share BLAST bit scores > 100. Colored nodes contain at least one sequence found in SwissProt, CARD, and BLDB, while the grey nodes do not have any known or characterized sequences. The size of the nodes indicates the number of sequences for each node in log10 scale. **b**, Distribution of protein families within the PBP-like superfamily. Categorization of various protein families within the PBP-like superfamily. The total number of sequences for each subfamily is determined by retrieving sequences from meta-nodes containing experimentally validated sequences and neighbouring connected meta-nodes. **c**, separation of SBLs classes in PBP-like superfamily.Summary of the SBL classes separation bit score cutoff within the PBP-like superfamily. The bottom axis indicates the SSN clustering threshold defining each class of SBL. The SSN nodes are shown at every 10 bit score, and the exact position of each nodedoes not cover the clustering thresholds in-between bit score intervals.

**Supplementary Fig. 3.**
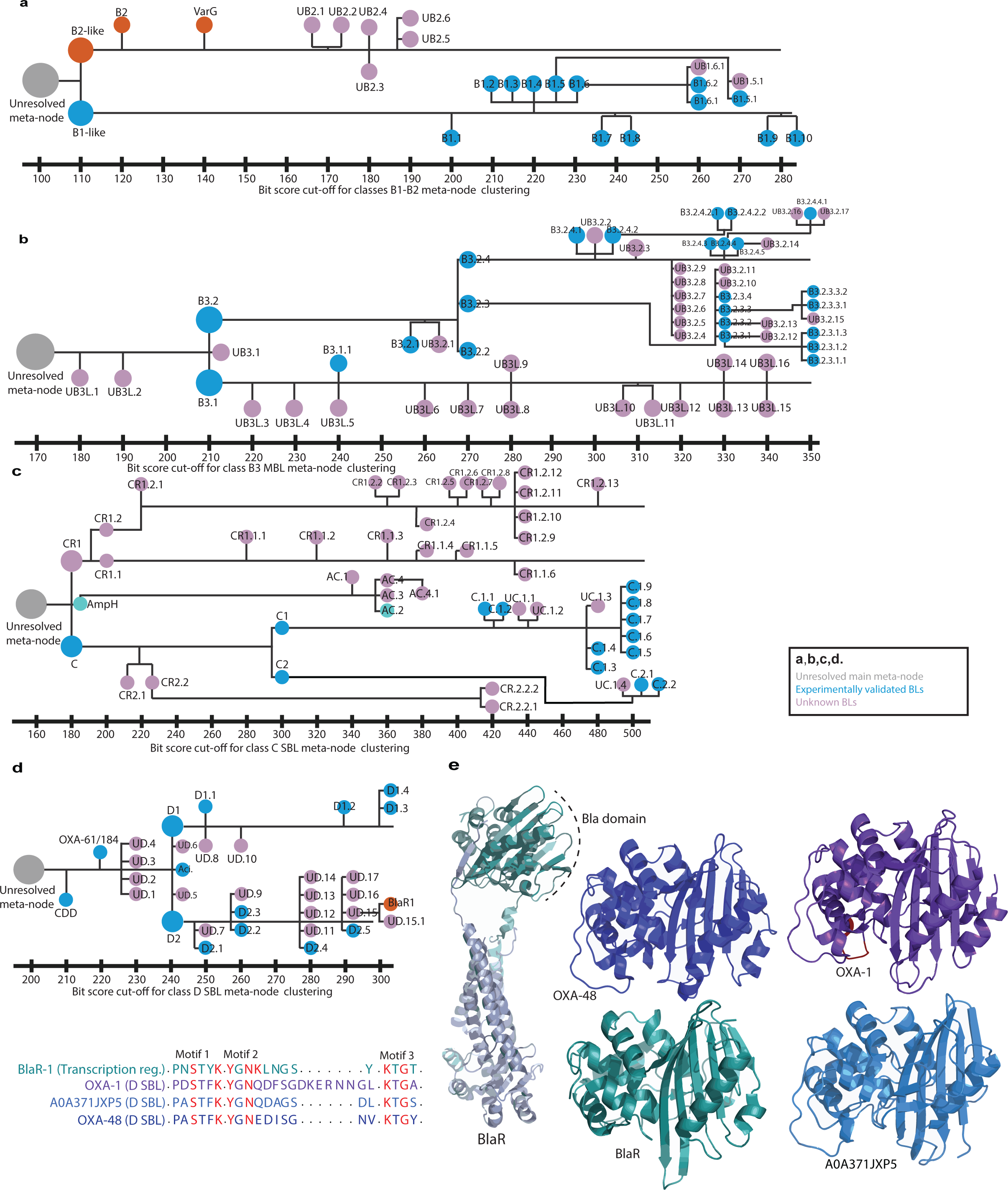
Overview of β-lactamase classes seperation. Summary of the BL classes seperation. Summary of the separation of meta-nodes (with more than 100 sequences) within the classes B1-B2 MBLs (**a**), B3 MBLs (**b**), C SBLs (**c**), D SBLs (**d**). In figuers a-e, SSN nodes are shown at every 10 bit score, and the exact position of each node does not cover the clustering thresholds in-between bit score intervals. The dark blue nodes contain at least one β-lactamase sequence found in CARD and BLDB, while the light pink nodes (UA, UC, UD, UB) do not have any known or characterized β-lactamase sequences. The orange nodes contains B2,VarG and BlaR1 proteins while cyan and orange nodes contains AmpH LMW-PBP. UB1: Unknown class B1; UB2: Unknown class B2; UB3: Unknown class B3; UA: Unknown class A; UC: Unknown class C; AC: AmpH cluster; CR: class C related; UD: Unknown class D; Aci: Acinetobacter cluster. (**e**), Multiple sequence alignment and structures of representative class D SBLs.The highly consereved amino acids in active site motifs are highlited in orange. Overall structure from the class D β-lactamases and BlaR (OXA-1 (D), purple; OXA-48 (D), p-urple blue; A0A371JXP5 (D), light blue; BlaR (transcription regulator), green.

**Supplementary Fig. 4.**
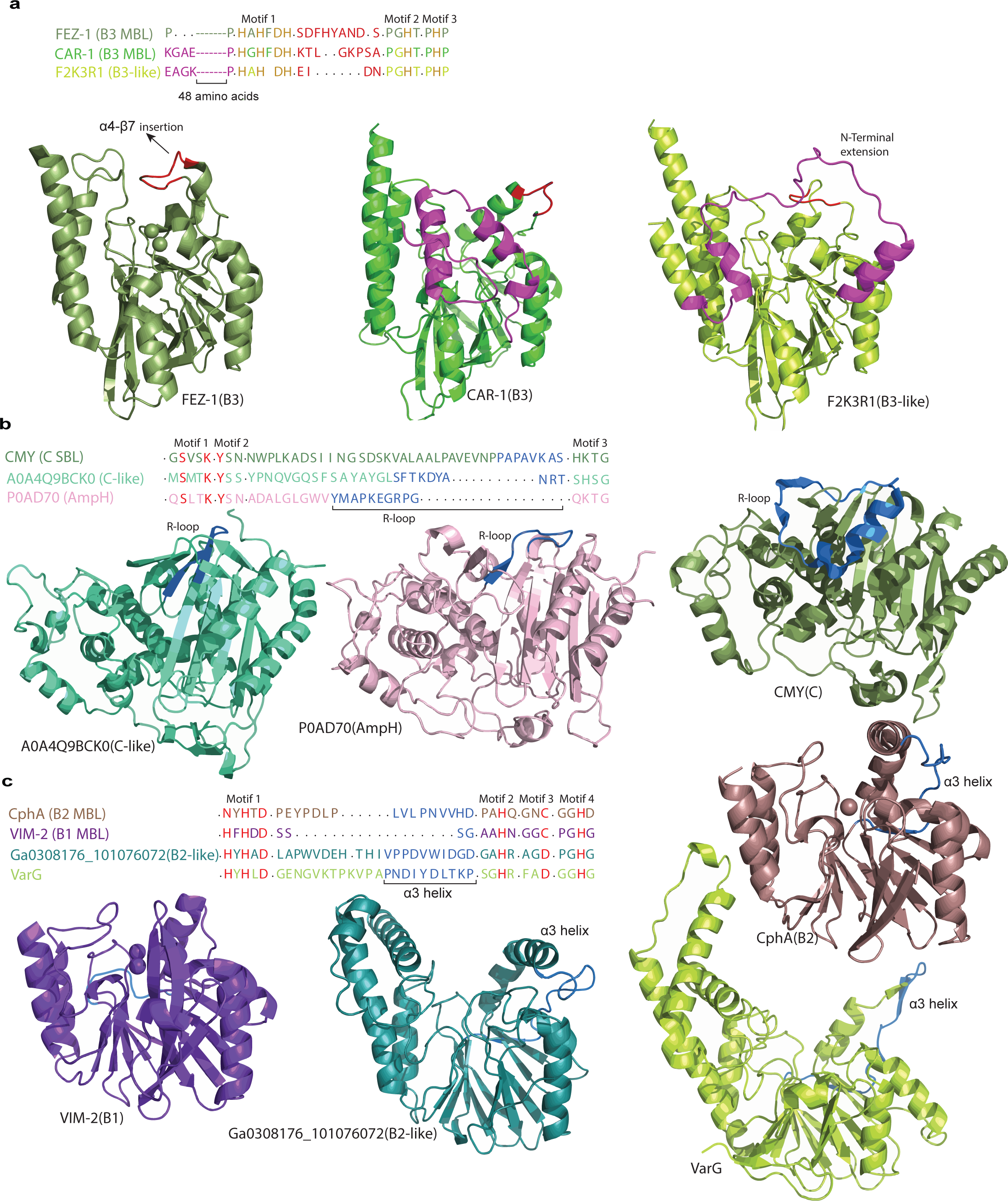
Multiple sequnce alignemnt and structures of the β-lactamase classes. **a**, Multiple sequence alignment and structures of representative class B3 MBLs.The highly consereved amino acids in active site motifs are highlited in orange. The N-Terminal B3-like and the loop insertion in between α4-β7 in B3 are colored in magneta and red. Overall structure from the class B3 and B3-like β-lactamases (FEZ-1 (B3), dark green; CAR-1 (B3), green; F2K3R1 (B3-like), light green). **b**, Multiple sequence alignment and structures of representative class C SBLs and ne--ighboring C-like sequences. The highly consereved amino acids in active site motifs and the R-loop are highlited in red and blue. overall structure and R-loop of re--presentatives from the class C and C-like β-lactamases (CMY (C), sage green; A0A4Q9BCK0(c-like), green cyan; P0AD70 (AmpH), light pink). **c**, Multiple sequen--ce alignment and structures of representative class B1-B2 MBLs.The highly consereved amino acids in active site motifs and the α3 helix are hig-hlited in red and blue. overall structure from the class B1-B2 and B-like sequences (VIM-2 (B1), purple; CphA (B2), brown; Ga0308176_101076072 (B2-like), turquoise; VarG, light green).

**Supplementary Fig. 5.**
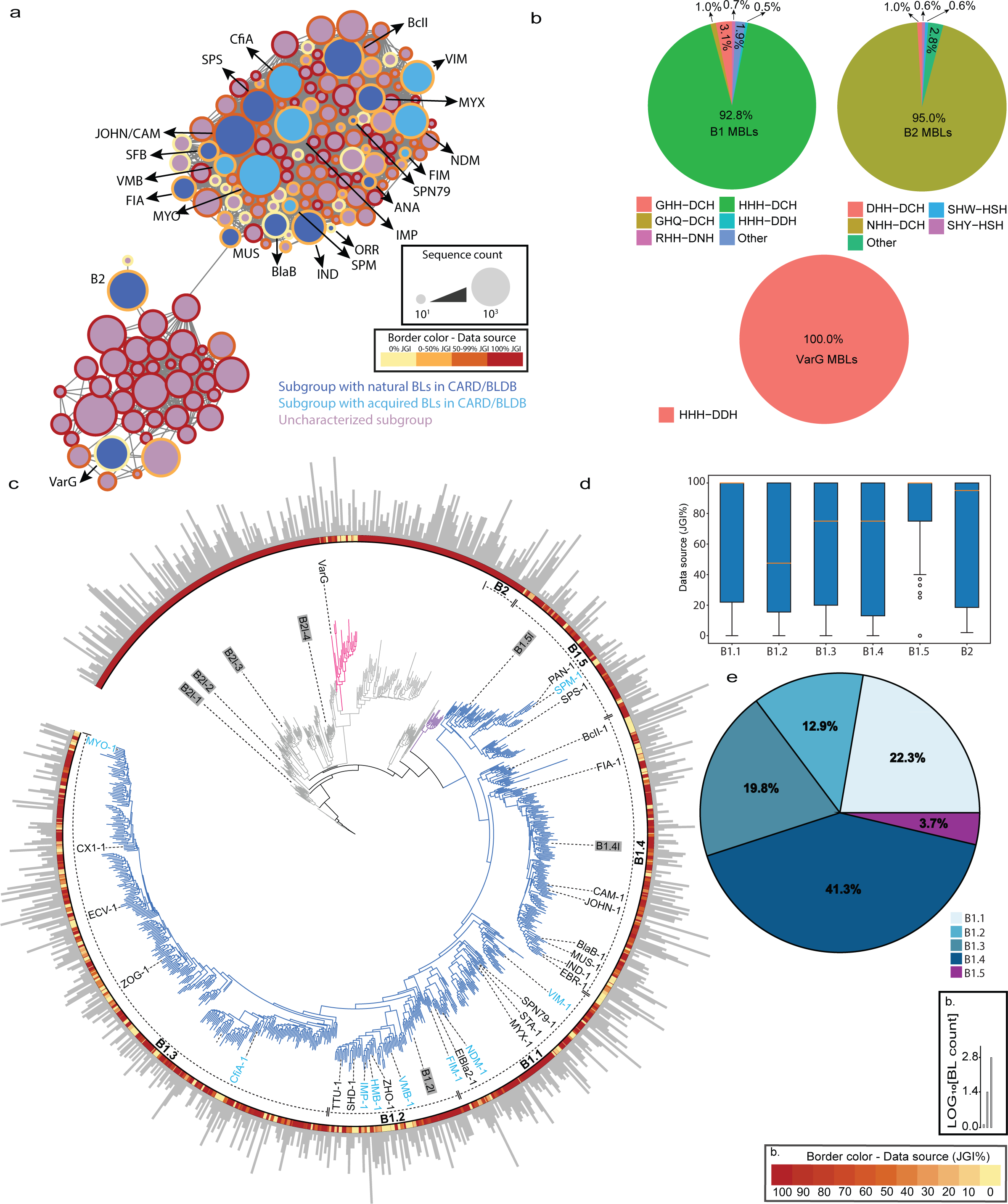
Sequence relationships among classes B1 and B2 metallo-β-lactamases (MBL) found in genomic and metagenomics databases. **a**, Sequence similarity network of class B1/B2 sequences. Nodes indicate groups of MBL sequences sharing BLAST bit scores > 280. Nodes are linked by edges if sequences in two separate nodes share BLAST bit scores > 50. Blue nodes contain at least one sequence found in CARD/BLDB, while the grey nodes do not have any known MBL sequences. The size of the nodes indicates the number of sequences for each node in the log10 scale. The node border color indicates the sequen--ce source composition. **b**, Distribution of active site motif within the B1, B2 and VarG MBLs. **c**, Phylogeny of representative sequences from class B1-B2 MBLs.Se--quences represent protein clusters with at least 60% sequence pairwise identities along with specific sequences of interest, with light blue names representing mo--bilized B1-B2 sequences. The branch colors show the percentage of the environmental sequence each phylogeny sequence represents.The inner dash lines indic--ate the B1 MBL clade names, while the sequence line color represents the MBL class. The outer bar plot illustrates the number of sequences each node represents. The experimentally characterized sequences are visually indicated by the colored box, with active proteins in blue and non-functional sequences in grey. **d**,Box-plot exhibiting the distribution of environemntal sequences in B1 clades. **e,** Pie-chart exhibiting the size of B1 clades.

**Supplementary Fig. 6.**
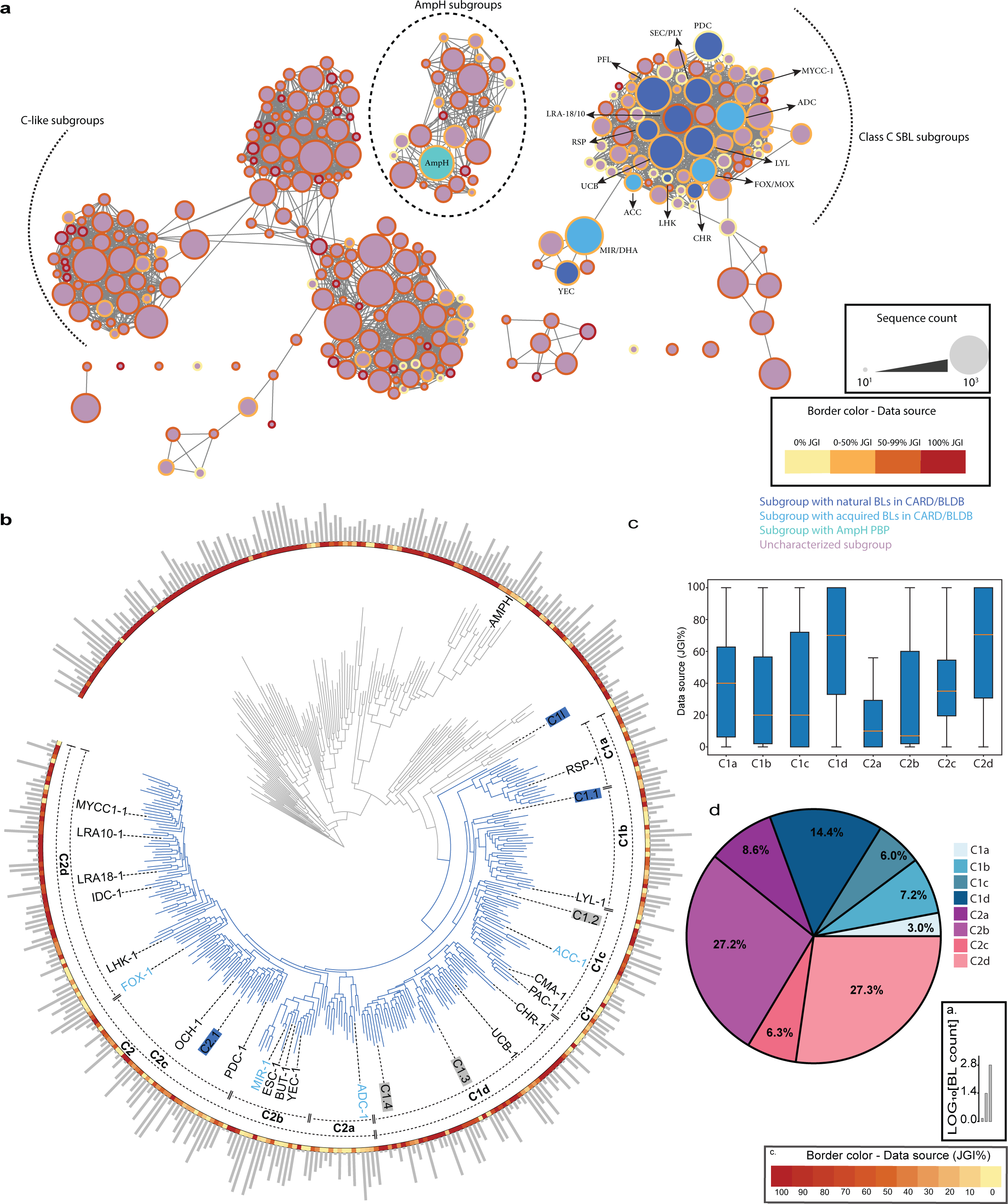
Sequence relationships among class C SBLs found in genomic and metagenomics databases. Summary of the sequence space for C SBLs. **a**, Sequence similarity network of class C sequences. Nodes indicate groups of MBL sequences (with more than five sequences) sharing BLAST bit scores > 500. Nodes are linked by edges if sequences in two separate nodes share BLAST bit scores > 300. Blue nodes contain at least one MBL sequence found in CARD and BLDB, while the grey nodes do not have any known or characterized SBL sequences. The size of the nodes indicates the number of sequences for each node in the log10 scale. The node border color indicates the sequence source composition for each node. **b**, Phylogeny of repr--esentative sequences from class C SBLs.Sequences represent protein clusters with at least 60% sequence pairwise identities along with specific sequences of int--erest (annotated), with light blue names representing mobilized class C sequences. The branch colors show the percentage of the environmental sequence each phylogeny sequence represents.The inner dash lines indicate the class C SBL clade names. The outer bar plot illustrates the number of sequences each node rep--resents.epresents.The experimentally characterized sequences are visually indicated by the colored box, with active proteins in blue and non-functional sequences in grey. **c**,Box-plot exhibiting the distribution of environemntal sequences in C clades. **d,** Pie-chart exhibiting the size of C clades.

**Supplementary Fig. 7.**
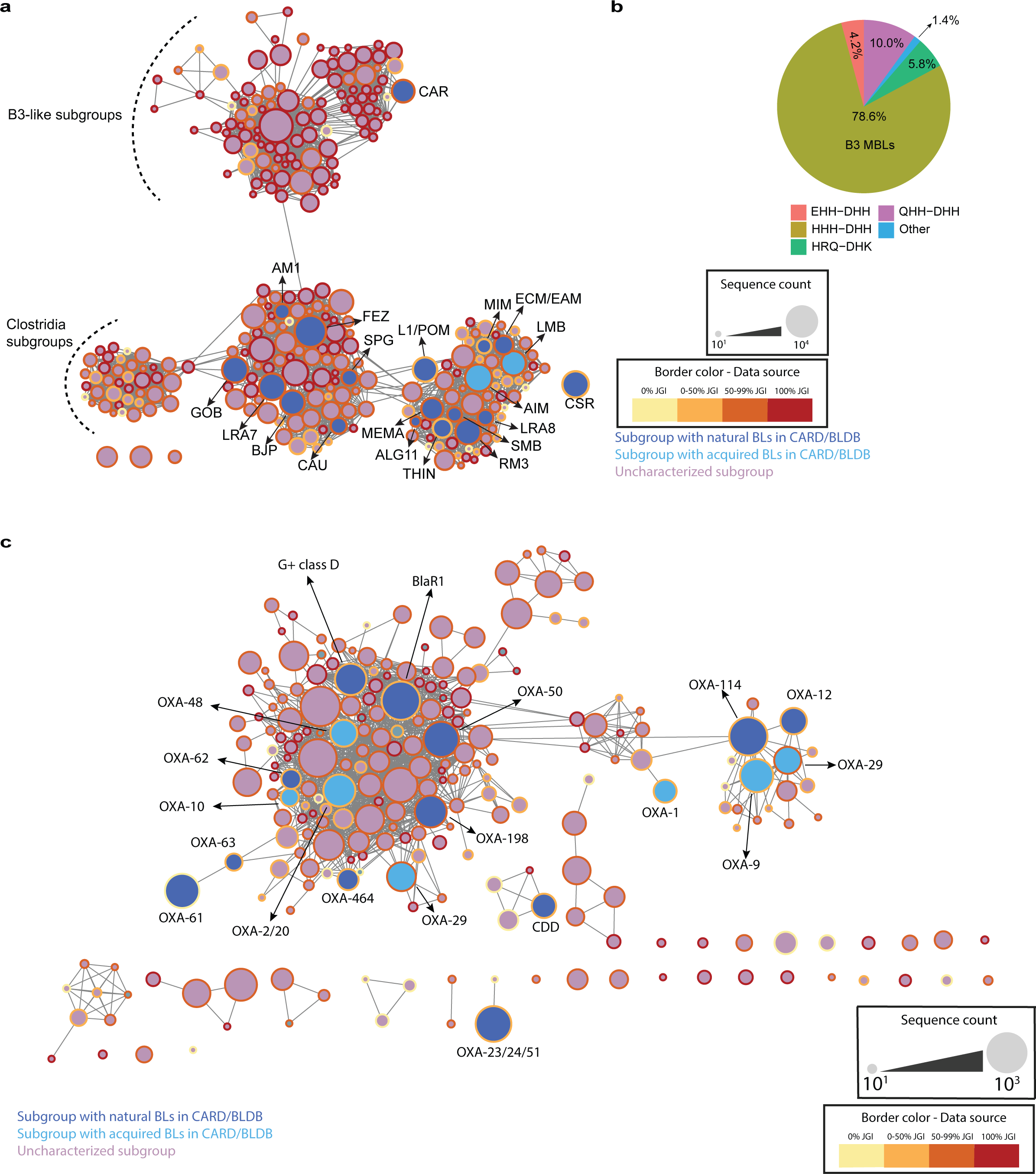
Sequence relationships among classes B3 MBLs and D SBLs found in genomic and metagenomics databases. Summary of the sequence space for B3 and D BLs. **a**, Sequence similarity network of class B3 sequences. Nodes indicate groups of MBL sequences sharing BLA--ST bit scores > 350. Nodes are linked by edges if sequences in two separate nodes share BLAST bit scores > 160. **b**, Distribution of active site motif within the B3 MBLs. **c**,Sequence similarity network of class D sequences. Nodes indicate groups of SBL sequences (with more than five sequences) sharing BLAST bit scores > 300. Nodes are linked by edges if SBLs in two separate nodes share BLAST bit scores > 210. In both SSNs, Blue nodes contain at least one SBL sequence found in CARD and BLDB, while the grey nodes do not have any known or characterized SBL sequences. The size of the nodes indicates the number of sequences for each node in log10 scale. The node border color indicates the sequence source composition for each node.

**Supplementary Fig. 8.**
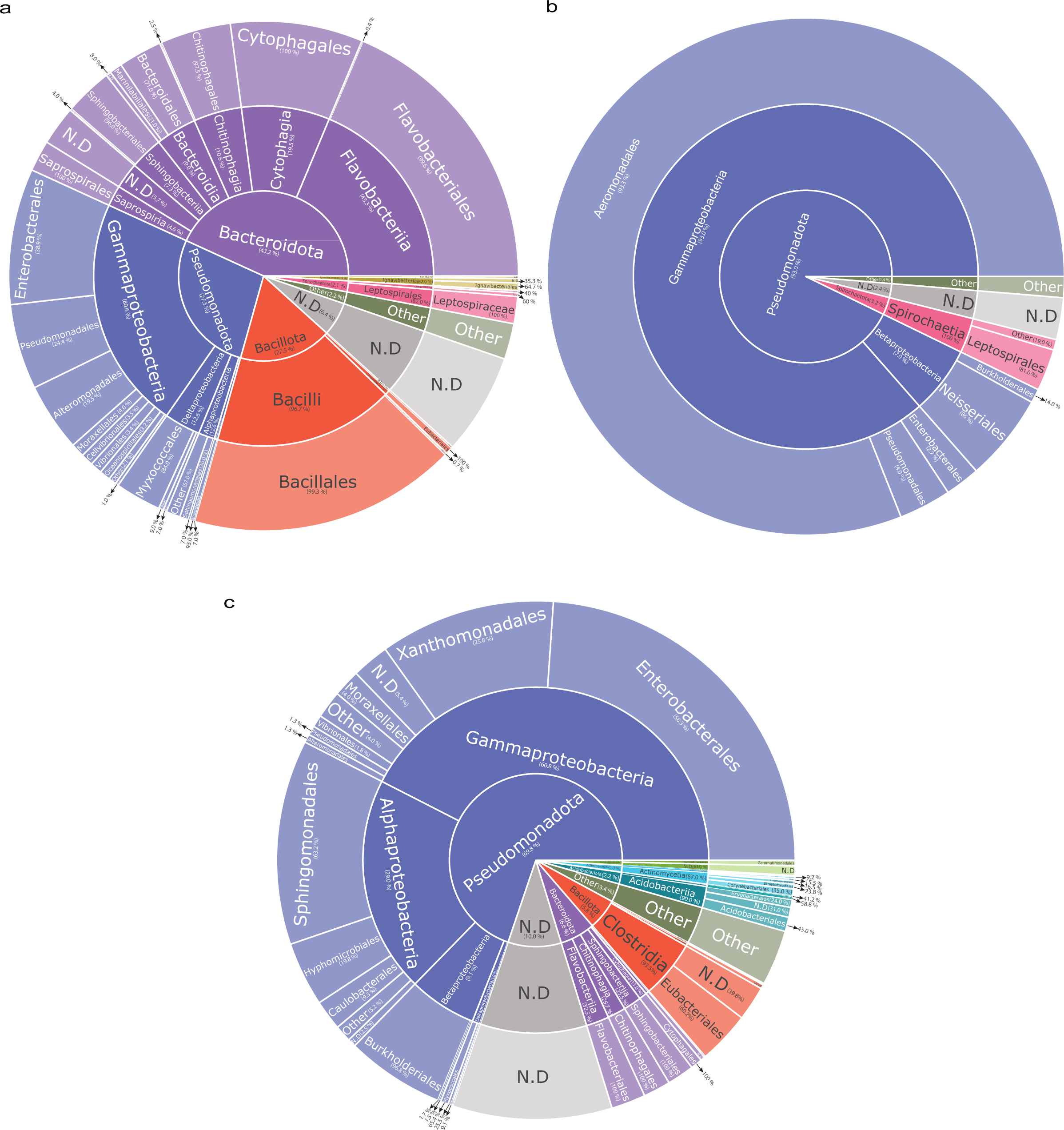
Taxonomic distribution of putative MBL sequences. Multi-level pie-chart of the taxonomic distribution of all classes of MBLs across phylum, class, and order taxonomic ranks. **a**, The taxonomic distribution of putative identifed sequecnes for classes B1 (**a**), B2 (**b**), and B3(**c**) MBLs.The sum of percentages for each layer is 100%.

**Supplementary Fig. 9.**
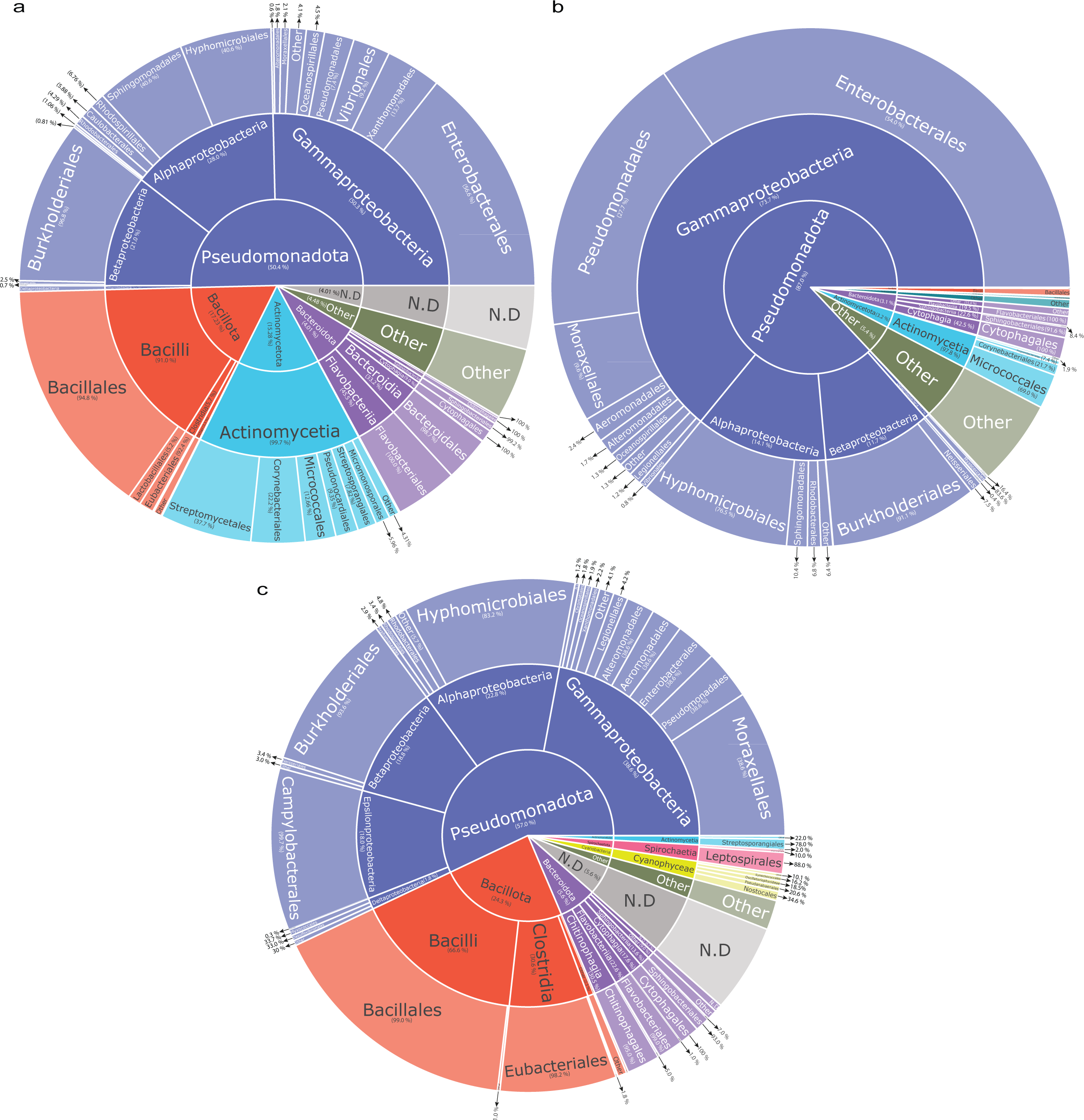
Taxonomic distribution of putative SBL sequences. Multi-level pie-chart of the taxonomic distribution of all classes of SBLs across phylum, class, and order taxonomic ranks. **a**, The taxonomic distribution of putative identifed sequecnes for classes A (**a**), C (**b**), and D(**c**) SBLs.The sum of percentages for each layer is 100%.

**Supplementary Fig. 10.**
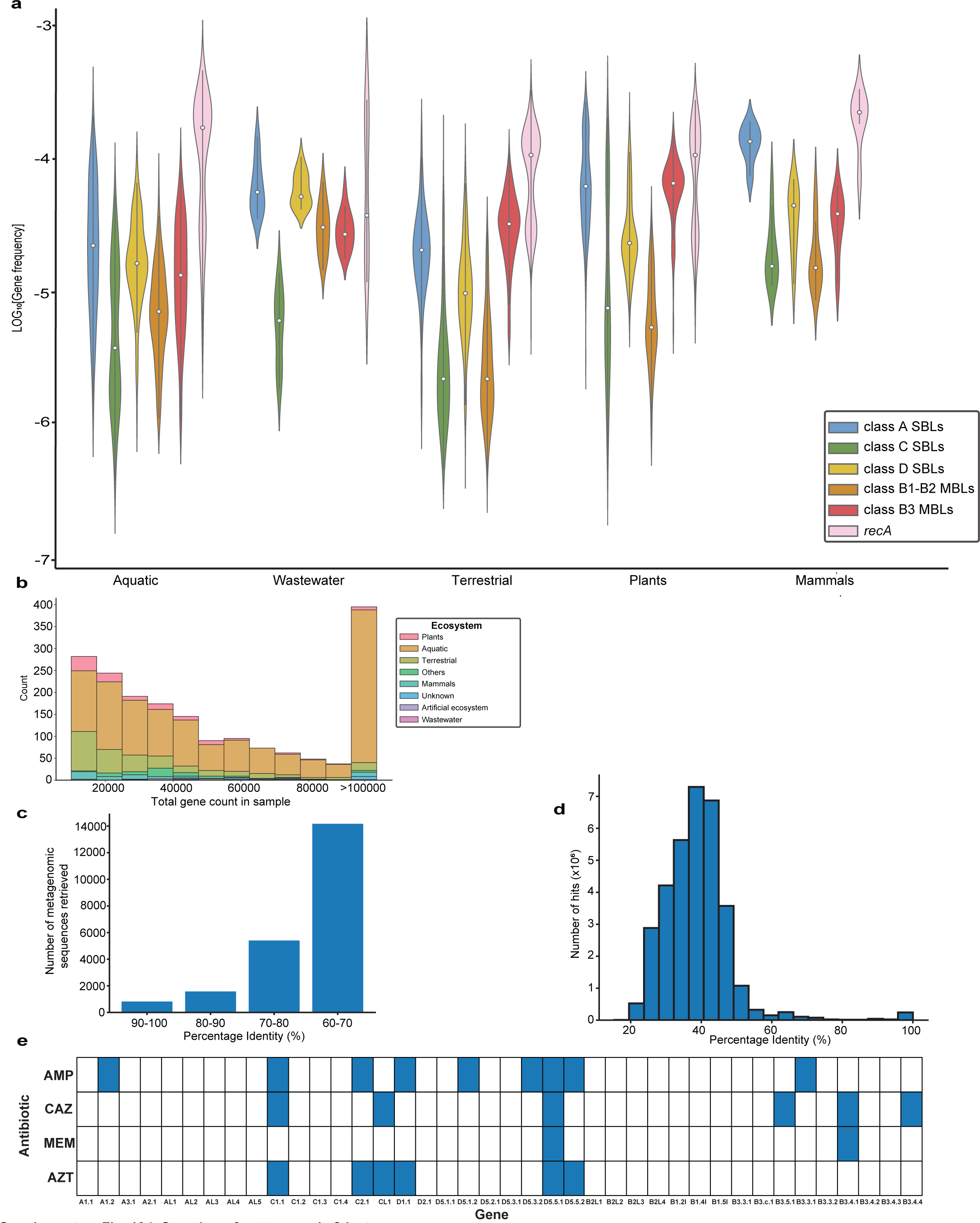
Overview of metagenomic β-lactamases. **a**, Violin plots showing the diversity of all classes of β-lactamases’ relative frequency in different ecosystems. The distribution of the *recA* marker gene is shown in pi--nk. **b**, Summary of samples with no identifed *bla* but at least one *recA* gene. **c**,The distribution of the average alignment identity, as measured by BLAST, of all pair--wise sequence comparisons of the identified metagenomic sequences against the sequences within the BLDB/CARD database. **d**, The distribution of the average alignment identity of all pairwise sequence comparisons of the identified sequences against the Resfinderfg v4.0 database. **e**, Heatmap showing the antibiotic resist--ance phenotype for 40 sampled metagenomic sequences. Sequences ending in L (i.e., CL, DL, and B1L) are those predicted to be not β-lactamases (AMP: ampici--llin; CAZ: ceftazidime; MEM: meropenem; AZT: aztreonam).

